# Motor behaviour selectively inhibits hair cells activated by forward motion in the lateral line of Zebrafish

**DOI:** 10.1101/772129

**Authors:** Paul Pichler, Leon Lagnado

## Abstract

How do sensory systems disambiguate events in the external world from signals generated by the motor behaviour of the animal? One strategy is to suppress the sensory input whenever the motor system is active, but the cellular mechanisms remain unclear. We investigated how motor behaviour modulates signals transmitted by the lateral line of zebrafish, which senses pressure changes around the body of the animal. Activation of motor neurons during fictive swimming caused co-activation of efferent fibers and suppression of synaptic transmission from the primary mechanoreceptors, the hair cells. In some hair cells, a single motor spike inhibited glutamate release by about 50% and block was often complete within 50-100 ms of the start of swimming. All hair cells polarized to be activated by posterior deflections, as would occur during forward swimming, were suppressed by >90%, while only half of those polarized in the anterior direction were inhibited and by an average of just 45%. The selective inhibition of hair cells activated during motor behaviour provides a mechanism for the suppression of self-generated signals while maintaining sensitivity to stimuli originating in the external world.

## Introduction

Animals need to distinguish sensory signals originating in their environment from signals generated by their own behaviour within that environment. During vision, for example, backwards-drifting optic flow can be the result of active forward locomotion or the animal’s displacement caused by external forces such as wind or water flow. These two situations are handled very differently: when a fish or a fly experiences passive optic flow the optomotor response is triggered to stabilize its position relative to the visual world, but when the animal actively engages motor systems to move forward this innate behaviour becomes counterproductive and is therefore suppressed. Such a behavioral switch could in principle be generated if a copy of the motor command was transmitted to the sensory apparatus to alter processing of the self-induced sensory signals and strong evidence for this has recently emerged in the visual system of Drosophila (Erich and Horst, 1950; Kim et al., 2017; Kim et al., 2015). But the cellular mechanisms by which an efference copy acts on the sensory apparatus are still unclear.

A second sensory system in which active locomotion must be disambiguated from external events is the lateral line of fish and amphibians. The lateral line senses pressure changes in the water around the body and is involved in a number of behaviors and reflexes, such as obstacle and predator avoidance and startle and escape responses (Burgess and Granato, 2007; Haehnel-Taguchi et al., 2014; Troconis et al., 2017). Of particular importance is rheotaxis, whereby an aquatic animal stabilizes itself in the face of water flow by making compensatory swimming motions (McHenry et al., 2009; Stewart et al., 2013). In larval zebrafish, rheotaxis can occur in the dark, driven by the lateral line detecting flow velocity gradients around the body (Oteiza et al., 2017). Forward swimming motions required for rheotaxis will, however, themselves activate the lateral line. How effectively are such re-afferent signals prevented from triggering a reflex? And does this come at the cost of also blocking transmission of external signals?

The end organs of the lateral line, the neuromasts, are distributed over the body surface. Each contains ~15-20 hair cells projecting their ciliary bundles into a single structure, the cupula. Hair cells are polarized to be activated either by posterior deflection of the cupula, as occurs during swimming, or by anterior deflection: signals transmitted from hair cells of each polarity are segregated through afferent fibres selective for that polarity. Neuromasts also receive inputs from cholinergic efferents that modulate the sensory signal transmitted to the hindbrain (Bricaud et al., 2001; Chagnaud et al., 2015; Flock and Russell, 1976). The mechanical sensitivity of hair cells varies, but some detect displacements of just a few tens of nanometers (Pichler and Lagnado, 2019).

Here we investigate how motor behaviour modulates the transmission of mechanical information in the lateral line of larval zebrafish by *in vivo* imaging of both neural and synaptic activity. We demonstrate that efferent neurons transmit a precise copy of the motor signal to the neuromast to modulate glutamate release from hair cells and that the sensitivity of the modulatory system is such that a single spike in the motor nerve inhibits release by ~50% within 50 ms. The efference copy signal does not, however, act uniformly within the neuromast: hair cells polarized to be activated by posterior deflections, as would occur during forward swimming, are suppressed by >90% while only half of those polarized in the anterior direction are inhibited and by an average of just 45%. These results demonstrate that the efference copy can completely suppress re-afferent signals generated by motor behaviour while preserving some sensitivity to stimuli originating in the external world.

## Results

### Neuromasts receive an almost exact copy of the motor signal

Cholinergic efferents entering neuromasts are thought to be co-activated with motor neurons to provide feedforward control of the sensitivity of the lateral line (Chagnaud et al., 2015) but the quantitative relationship between motor activity and the efferent and afferent signals are not known. To under these aspects of the systems operation, we made simultaneous measurements of activity in the motor nerve and efferent neurons and then related these to the stimulus-evoked output from hair cells as well as responses transmitted by afferent neurons. We used an *in vivo* preparation of transgenic zebrafish larvae (5-9 dpf) that undergo fictive swimming while neuromuscular transmission is blocked (Masino and Fetcho, 2005). Motor nerve activity was measured electrophysiologically while optical reporters were targeted genetically to measure calcium activity in efferent and/or afferent neurons as well as glutamate release from the synapses of hair cells in neuromasts towards the back of the tail (the posterior lateral line). These various measurements were combined with the application of mechanical stimuli to individual neuromasts to assess changes in sensitivity (**Fig 1A – D**; Pichler and Lagnado, 2019).

**Figure 1.**
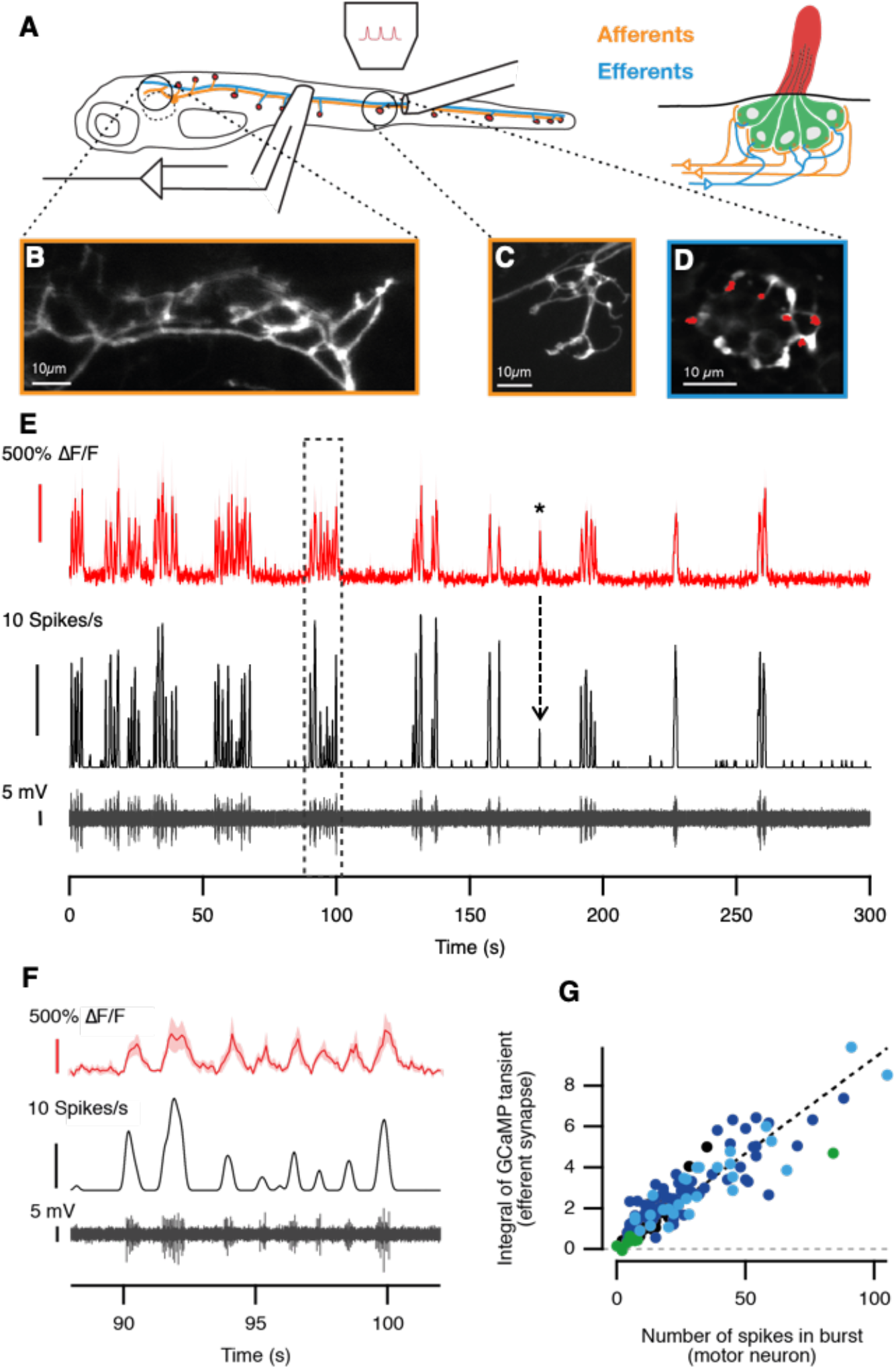
The efferent signal is an almost exact copy of the motor signal during fictive swimming. (**A**) At 7 dpf, the posterior lateral line of larval zebrafish consists of 14 neuromasts on each side (red dots). Each neuromast is innervated by at least two afferent neurons (yellow) and an efferent (blue). We imaged glutamate release of individual hair cells in a neuromasts while measuring motor neuron activity through a suction pipette. A second pipette applied pressure steps to the neuromast. (**B**, **C**) Average projections of the afferent synapses in the hindbrain (B), and a neuromast (C) of a larva expressing iGluSnFR under transcriptional control of the SILL promoter (Tg[Sill2, UAS::iGluSnFR]). (**D**) Average projection of a neuromast in a larva expressing GCaMP6f under the transcriptional control of the HuC (elavl3) promoter (Tg[HuC::GCaMP6f]), in which afferents and efferents (but not hair cells) are labelled. Red dots indicate efferent ROIs identified based on their firing pattern. (**E**) Top trace (red): spontaneous activity of the synapses in (D) over a five-minute period without visual or mechanical stimulation. The lower traces (black) depict the raw motor activity and the spike rate. The asterisk indicates a signal in the efferent synapses that correlates to six spikes in the motor nerve. (**F**) Magnified view of the dashed area in (E), showing that efferent synapses in the neuromast are activated at each swim-bout. (**G**) The number of spikes per swimming bout and the integral of the fluorescent signal during that episode were strongly correlated (r=0.9, n=155 bouts from 4 NMs, each depicted in a different colour).

To monitor the efferent signal we used the elevl3::GCaMP6f line of fish that express the calcium indicator GCaMP6f in afferent and efferent fibres but not hair cells (Faucherre et al., 2009) **Fig 1D**). Pre-synaptic boutons of efferent fibres could be distinguished from the post-synaptic varicosities of afferent neurons both by their smaller and rounder shape (Nagiel et al., 2008); red regions in **Fig 1D**) and by the effects of a mechanical stimulus – afferents were excited while efferents were not affected (**Fig S1**). In all 15 neuromasts tested, fictive swimming caused efferent synapses to be activated in a burst-like fashion in close synchrony with the spiking activity of the motor nerve (**Fig 1E**). Activity in the motor nerve and efferent fibre was tightly coupled: each burst of spikes in the motor nerve was associated with a calcium transient in efferent synapses (**Fig 1F**) and the number of spikes in a bout was directly proportional to the time-integral of the signal from GCaMP6f (**Fig 1G**). As few as 6 spikes within a motor burst was sufficient to generate a sizeable calcium signal in the efferent synapses (asterisk in **Fig 1E**). These results demonstrate that the efference copy signal transmitted to the neuromast copies the motor signal driving locomotion both quantitatively and temporally. Further, activity across all the efferent synapses within a single field of view were closely synchronized indicating that all hair cells within the neuromast receive a similar modulatory signal irrespective of their polarity (**Fig S2**).

### The efference copy suppresses both spontaneous and stimulus-evoked transmission from hair cells

To what extent does the efference copy modulate the output from a neuromast? To investigate this question we monitored the synaptic output from hair cells by expressing the glutamate sensor iGluSnFR (Marvin et al., 2013) under the control of the *Sill* promoter (Pichler and Lagnado, 2019; Pujol-Marti and Lopez-Schier, 2013); **Fig 1B & C** and **Fig 2A & B**). In these experiments we did not paralyze fish by the usual method of applying the neuromuscular blocker α-BTX because this agent has also been reported to block the α9/α10 isoforms of nicotinic acetylcholine receptors (nAChR) present in hair cells (Erickson and Nicolson, 2015; Verbitsky et al., 2000). Instead, we expressed iGluSnFR in the background of the *relaxed* mutant (cacnb^ts25/ts25^) in which defective dihydropyridine receptors block excitation-contraction coupling in muscles (Bohm et al., 2016; Granato et al., 1996; Schredelseker et al., 2005).

**Figure 2.**
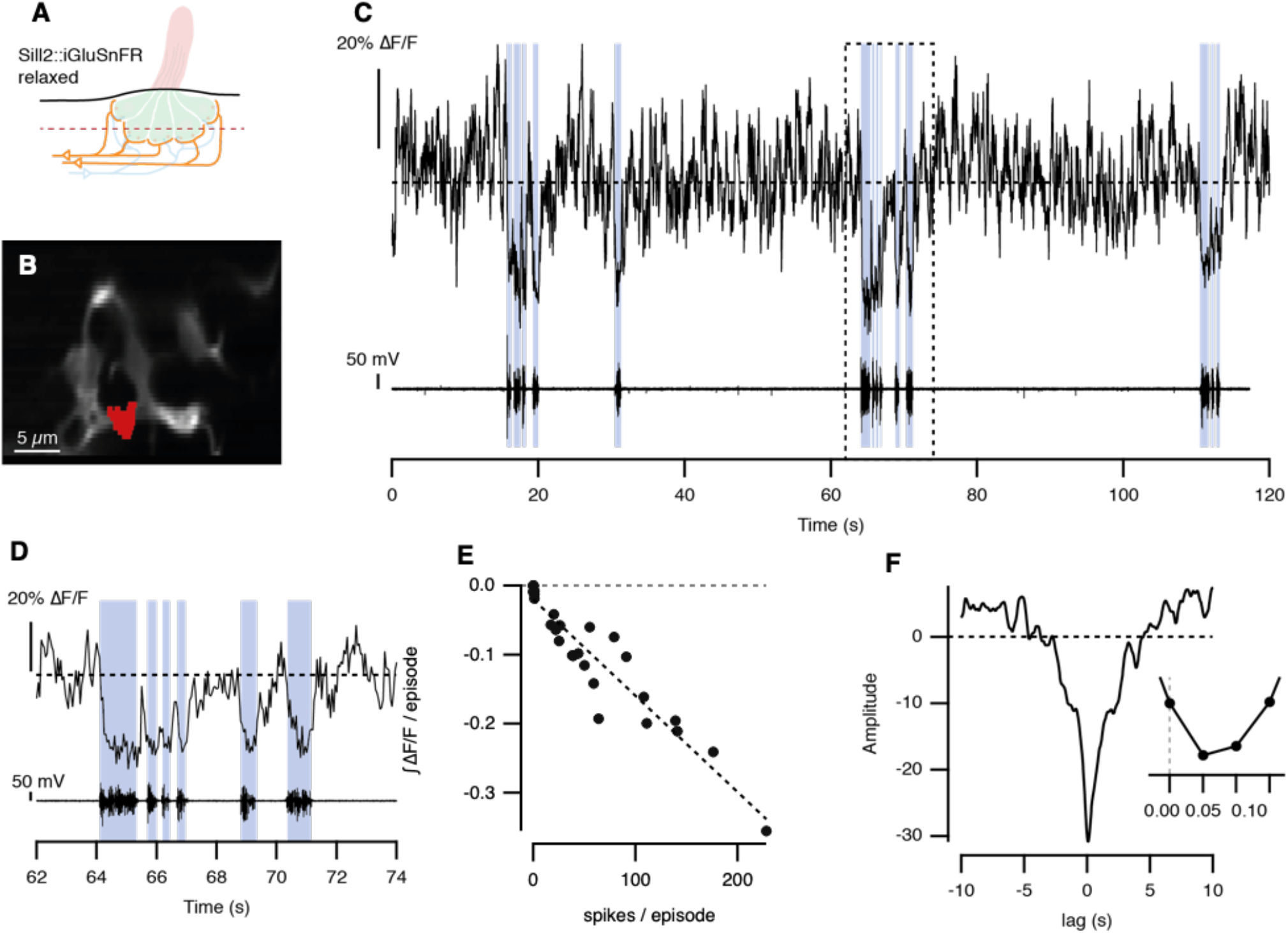
Spontaneous release of glutamate from hair cells is suppressed during fictive swimming. (**A**) Experiments were carried out in ‘relaxed’ mutants that express the glutamate reporter iGluSnFR in afferent neurons (Tg[Sill2, UAS::iGluSnFR], cacnb^ts25/ts25^) at 5 dpf. (**B**) A representative synapse, labelled in red, whose activity is presented in (C). (**C**) Spontaneous glutamate release of a hair cell over a period of two minutes (top panel) and motor neuron activity (lower panel). Blue areas indicate episodes of motor neuron activity. (**D**) Magnified view of boxed area in (C). The maximum suppression of glutamate release was similar for each burst of motor activity. (**E**) Relationship between the number of spikes in a burst and the negative integral of the iGluSnFR signal from the MON depicted in B (n = 28 swimming episodes, r = −0.95). (**F**) Cross-correlation of iGlusnFR signal and the number of spikes (downsampled to match imaging frequency). Inset shows that the peak is at 50 ms, the imaging interval.

Bouts of fictive swimming reduced the spontaneous release of glutamate from hair cells occurring in the absence of a stimulus, as shown by the example in **Fig 2C**. Suppression was evident for each burst of motor activity (**Fig 2D**), and the integral of the decrease in the iGluSnFR signal during a burst was directly proportional to the number of spikes it contained (**Fig 2E**). Cross-correlating the iGluSnFR signal with the motor nerve recording revealed that suppression was maximal within 50 ms of a spike, which was within the temporal resolution of image acquisition (**Fig 2F**). The suppressive effect of the efference signal was rapidly reversible: spontaneous release of glutamate from the hair cell recovered within ~100 ms of the end of a burst of spikes (**Fig 2D**). Similar suppression of spontaneous activity was observed in four neuromasts out of eight, and could also be observed by monitoring glutamate release at the output of afferent neurons in the Medial Octavolateralis Nucleus (MON) (**Fig S3**). Synaptic activity in the absence of a mechanical stimulus is a key aspect of the “push-pull” system by which a population of hair cells of opposite polarity signals stimulus direction (Pichler and Lagnado, 2019) and blocking spontaneous release will prevent the signalling of direction by the hair cells *inhibited* by a particular direction of motion.

Stimulus-evoked release of glutamate from hair cells was also suppressed during fictive swimming. In these experiments, we stimulated individual neuromasts with positive and negative pressure steps that deflected the cupula along the anterior-posterior axis while simultaneously measuring motor nerve activity and synaptic release of glutamate onto afferent neurons (**Fig 3**). These pressure steps were sufficient to generate maximal responses, and examples of glutamate signals from two neuromasts are shown in **Figure 3A and B** and **Figure 3D and E**, respectively. In each case we show signals from two hair cells, one polarised to be excited by posterior deflections of the cupula (green traces) and the other to anterior deflections (red traces). In neuromast 1, the hair cell signalling posterior deflections was markedly suppressed whenever mechanical stimulation overlapped with periods of motor nerve activity (**Fig. 3B**, highlighted in blue). In contrast, the hair cell signalling anterior deflections was unaffected. Responses in the boxed areas ‘1’ and ‘2’ are shown on an expanded time-scale in **Fig 3E (left)**, where they have been superimposed on the average response of the same synapse in the absence of fictive swimming (dashed lines). In the affected hair cell, the iGluSnFR signal was strongly reduced within 50 ms of the onset of motor activity.

**Figure 3.**
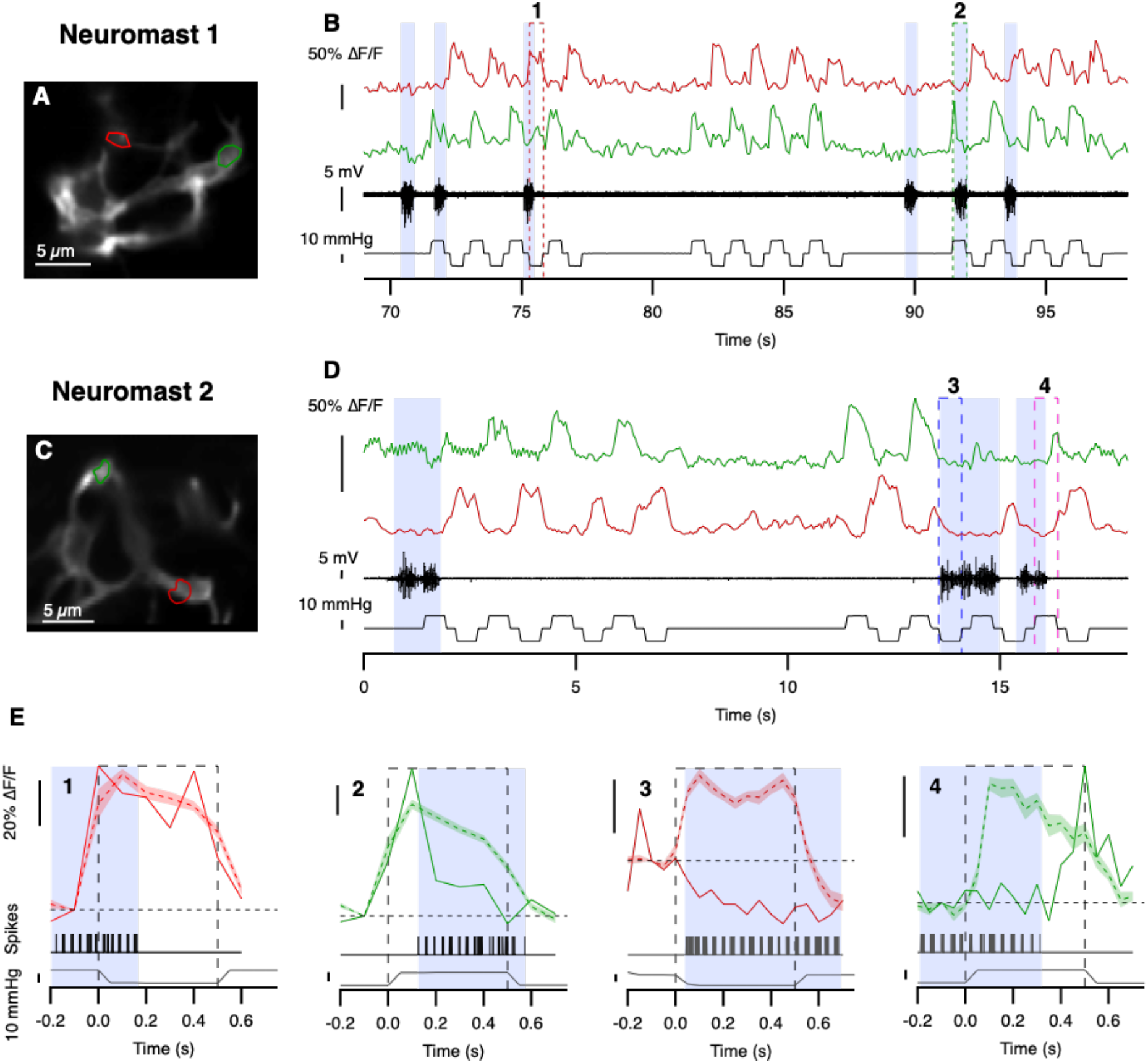
Motor behaviour blocks synaptic transmission from a subset of hair cells. (**A and B**) Image of iGluSnFR expression in afferents of neuromast 1 (A). Two representative synaptic inputs are highlighted in red (activated by anterior deflection) and green (activated by posterior deflection). The responses of these synapses to mechanical stimuli are shown in (B), together with motor nerve activity (black traces) and pressure steps applied to the neuromast. Positive pressure steps correspond to posterior deflections of the cupula and negative steps to anterior deflections. Blue shading indicates periods of motor nerve activity and numbered boxes indicate the stimulation episodes that are magnified in (E). (**C and D**) A corresponding representation of hair cell activity in neuromast 2. (**E**) Expansion of records in boxes 1-4 in B and D. The superimposed dashed red and green traces indicate the average mechanically-induced response of that synapse in the absence of motor nerve activity. Shaded areas represent the SEM. In example 2, inhibition of glutamate release is almost complete within 50 ms of the beginning of the motor burst. In example 3 suppression is complete within 50 ms, and further motor activity reduces glutamate release below resting levels. In example 4, glutamate release begins to recover within 50 ms of the end of the motor burst.

In neuromast 2, the two hair cells polarized to anterior and posterior deflections were *both* suppressed during fictive swimming (**Fig 3 D, and boxed areas ‘3’ and ‘4’ in E**). A particularly profound reduction in gain is evident in the examples highlighted in box 3, where the response to the mechanical stimulus was not simply nulled: glutamate release fell *below* the pre-stimulus baseline indicating that the efference copy signal was strong enough to also block a relatively high rate of spontaneous synaptic activity. Again, the suppressive effect of the efference copy signal could also be observed at later stages of signal transmission through the lateral line: glutamatergic output from afferent projections to the MON was strongly suppressed (**Fig S4**).

We surveyed the effects of motor activity on the output from 41 hair cell synapses from 8 neuromasts in 6 fish, and found that in 29 (71%) the response to a strong mechanical stimulus was significantly suppressed (**Fig. 4**). This assessment began by quantifying the relationship between motor activity and glutamate release using a metric, the Suppression Index (SI), that was calculated for each application of a mechanical stimulus. If Ro(*t*) is the average iGluSnFR signal at time *t* in the *absence* of motor activity and Rm(*t*) is the signal during a single stimulus trial, then SI at each time t during the trial was calculated as

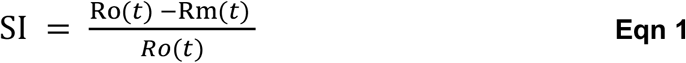

**Figure 4.**
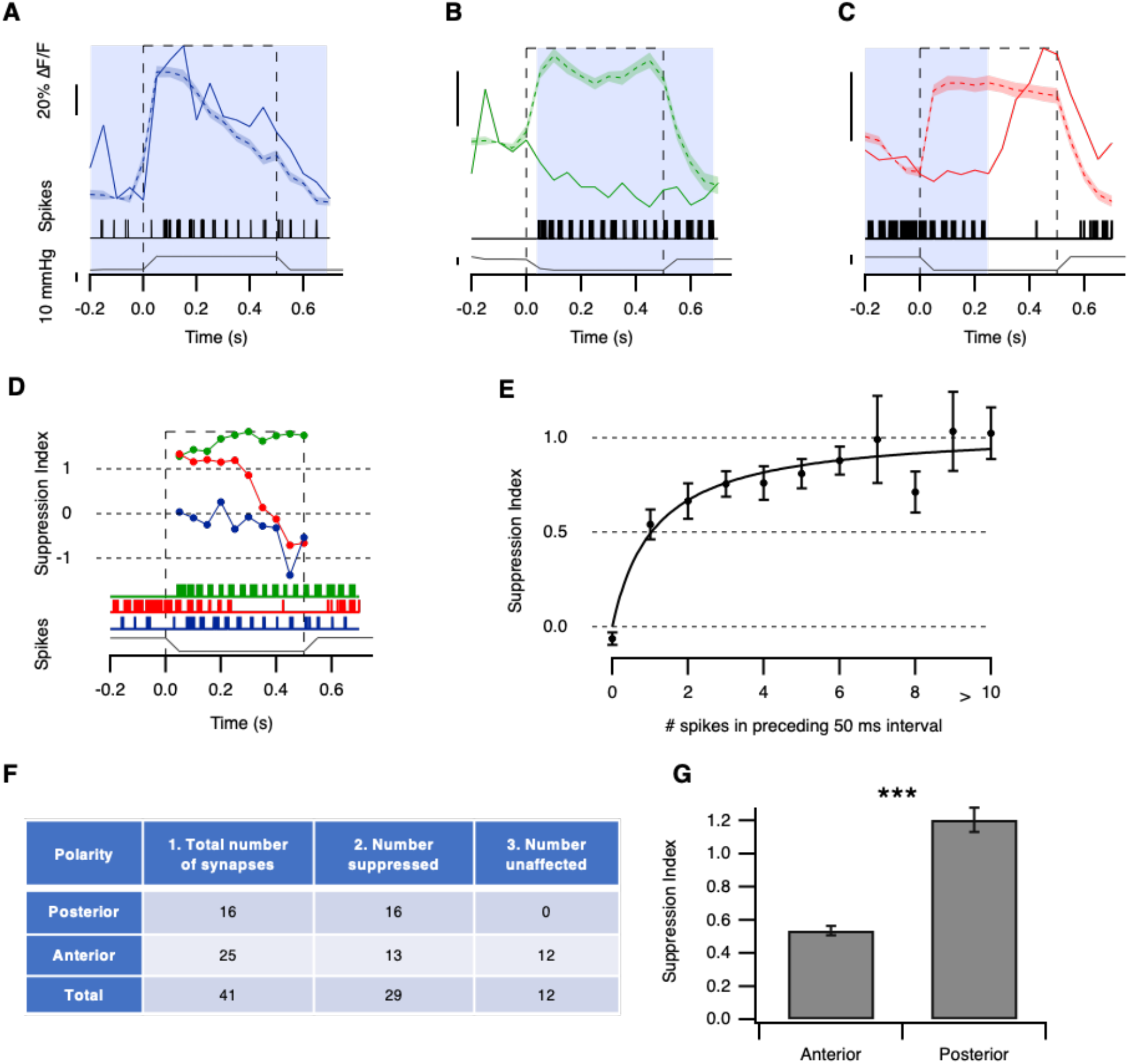
Motor behaviour selectively modulates hair cells activated by deflection in the posterior direction. (**A-C**) Three examples of synapses whose response was (A) unaffected, (B) suppressed during the entire stimulation episode and (C) suppressed only during the initial part of the stimulus. (Shaded areas represent the SEM.) (**D**) The suppression index (SI), calculated on a point-by-point basis during mechanical stimulation (equation 1). The red, green and blue traces show synapses from three different hair cells, with the corresponding motor activity shown below. Glutamate release from the blue synapse was not significantly suppressed (SI ~0); stimulated release from the red synapse was nulled during motor activity (SI ~1) but then recovered at the end of the burst of spikes; glutamate release from the green synapse was reduced to below resting levels (SI >1). (**E**) Plot of the relation between the SI at each time point during a mechanical stimulus and the number of spikes in the motor nerve in the preceding 50 ms time interval. Only synapses classified as suppressed were analyzed. Collected results from 29 synapses in 7 fish. The data could be described by a Hill equation of the form SI(N_s_) = (SI_max_*N_s_)/(N_s_ + N_1/2_) where N_s_ is the number of spikes, SI_max_ is the maximum SI (1.05 ± 0.08) and N_1/2_ is the number of spikes coinciding with half-maximal suppression (1.12 ± 0.42). Error bars show SEM. (**F**) The effects of motor activity on synaptic transmission from hair cells of opposing polarity. Column 1: The number of synapses activated by deflection in the posterior and anterior directions. Measurements were made in a total of 41 synapses in 8 neuromasts in 6 fish. Column 2: The number of synapses suppressed during motor activity, classified as described in the text. Column 3: The number of synapses unaffected by motor activity. Considering all synapses irrespective of polarity, the average probability of suppression was 29/41 = 0.7. Taking the null hypothesis as polarity having no bearing on suppression, the probability of observing suppression in all 16 synapses would be p = 0.7^16^ = 0.003. The null hypothesis can be rejected. (**G**) Comparison of the magnitude of the suppressive effect in hair cells polarized for anterior and posterior deflection. Only the 29 hair cells in which motor activity exerted a significant suppressive effect were analyzed. Suppression was stronger in posterior synapses (SI = 1.20 ± 0.03, n = 579 points) than in anterior synapses (SI = 0.54 ± 0.07, n = 709 points), a difference significant at p < 0.0001 using Mann-Whitney U test. Bars show SEM.

Thus, SI = 0 indicates no suppression, SI = 1 indicates full suppression and SI > 1 indicates suppression below the pre-stimulus baseline (i.e inhibition of glutamate release occuring at rest as well as complete nulling of the stimulus-evoked response). We only had limited control over the timing of fictive swimming relative to the application of the mechanical stimulus so SI could only be calculated when the two overlapped. Three examples of the calculation of SI are shown in **Fig. 4D**, based on responses shown in **Fig. 4A-C**. Each SI measurement over a 50 ms interval was then placed into one of two populations: intervals that coincided with at least one spike in the motor nerve and those that did not. These two populations were then compared using a one-tailed non-parametric Mann-Whitney U-test to assess whether motor activity had a significant effect on a stimulus-evoked response using a significance level of α=0.05. These results demonstrate that the efferent copy of the motor signal does not act uniformly on all hair cells within the neuromast.

Instantaneous spike rates of ~20 Hz in the motor nerve were sufficient to block the sensory signal transmitted by hair cells. This property of the efferent system was revealed by measuring the relation between the SI at each time point during a mechanical stimulus and the number of spikes in the preceding 50 ms time window. **Fig. 4G** shows collected results from 29 synapses in 7 fish. Half-maximal suppression was associated with an average of just 1.1 spike in the preceding 50 ms and 5 spikes were sufficient to cause an average of 80% suppression. Together, the results in Figures 1–4 demonstrate that motor activity acts rapidly and efficiently to block transmission of self-generated stimuli at the first synapse in the lateral line system.

### Efferent modulation is biased towards hair cells activated by posterior deflection

Ultrastructural studies indicate that *all* hair cells within a neuromast are innervated by efferent neurons (Dow et al 2018), but we found that motor activity only inhibited transmission from ~70% (**Fig. 4F**). To investigate this apparent discrepancy, we asked whether the effect of motor activity might depend on the polarity of the hair cell and found that it did: whereas 16/16 synapses polarized to posterior deflection were suppressed during motor activity, only 13/25 polarized to anterior deflection were affected (**Fig 4F**). Considering all synapses irrespective of polarity, the average probability of suppression was 29/41 = 0.7. Taking the null hypothesis as polarity having no bearing on suppression, the probability of observing suppression in all 16 synapses would be expected to be p = 0.7^16^ = 0.003: the null hypothesis can therefore be rejected. Further, of the 29 hair cells in which motor activity exerted a significant suppressive effect, the SI was greater in synapses activated by posterior deflection of the cupula (SI = 1.20 ± 0.03) compared to those activated by anterior deflection (**Fig. 4G**, SI = 0.54 ± 0.07; a difference significant at p < 0.0001 using Mann-Whitney U test). In other words, a burst of motor activity completely and selectively blocked transmission of a mechanical stimulus by the hair cells that would be most strongly activated by forward swimming motion while those of opposite polarity were still capable of signaling a stimulus.

### Discussion

Behavioural modulation of sensory information will necessarily involve a variety of different mechanisms, depending on the sensory modality (Chagnaud et al., 2015; Flock and Russell, 1976; Fujiwara et al., 2017; Keller et al., 2012; Kim et al., 2015; Saleem et al., 2013). Using the lateral line of zebrafish, we have found that efferents modulating the output of neuromasts are very tightly synchronized with motor activity (**Fig. 1**) and can rapidly and reversibly suppress transmission of mechanical signals from hair cells (**Figs. 2 and 3**) through to the central nucleus the MON (**Figs. S3 and S4**). Efferent modulation was strongly selective for hair cells activated by deflection towards the tail, and will therefore preferentially suppress self-generated signals occurring during forward swimming motion (**Fig. 4**).

The variable sensitivity of hair cells to the efference copy signal was unexpected because efferent fibres innervate all hair cells irrespective of their polarity (Faucherre et al., 2009). It appears that not all anatomical connections are equally effective in inhibiting hair cell activity. This might reflect presynaptic differences in the efficiency with which spikes trigger acetylcholine release and/or postsynaptic differences in the density of nicotinic receptors mediating calcium influx or calcium-activated potassium channels directly causing hyperpolarization (Dawkins et al., 2005).

Efferent modulation of hair cell activity will serve at least two functions. First, it will protect the lateral system from overstimulation (Roberts, 1989). In the absence of efferent modulation, swimming would generate a strong stimulus that would induce adaptation by, for instance, depletion of synaptic vesicles at the ribbon synapse of hair cells (Goutman, 2017; Pichler and Lagnado, 2019; Schnee et al., 2011; Schnee et al., 2005). At the ribbon synapse of auditory hair cells, recovery from synaptic depression is only complete after seconds of rest (Cho et al., 2011), which would introduce a “dead-time” at the end of a swimming bout when the lateral line was not signaling effectively. Secondly, the efference copy will prevent activation of reflexes normally triggered by the posterior lateral line, such as the escape response (Burgess and Granato, 2007; Troconis et al., 2017) or more gentle swimming triggered by stimulation of a single neuromast (Haehnel-Taguchi et al., 2014). These reflexes would cause positive feedback activation of motor activity if the efference copy signal did not break the loop by blocking the self-generated (re-afferent) signal. Blocking the lateral line system at source – the hair cells – will achieve this while preventing inappropriate activation of afferents or other downstream neurons.

The fact that only hair cells activated by anterior deflections of the cupula retained appreciable sensitivity does not mean that the lateral line system was unable to encode forward motion. We have previously demonstrated that hair cells within a neuromast form a heterogeneous population in terms of their sensitivity and set-point determining glutamate release at rest (Pichler and Lagnado, 2019). While some hair cells rectify completely and only encode deflections in their preferred orientation, others are able to encode opposing directions of motion by increasing or decreasing their glutamate release from a relatively high baseline. In the most extreme cases, ~40% of a hair cells dynamic range was occupied by deflections in their non-preferred direction. These cells would be able to encode forward swimming by *decreasing* glutamate release. Furthermore, some hair cells were found to generate a strong rebound release of glutamate on cessation of a stimulus in their nonpreferred orientation, encoding both its magnitude and duration (Pichler and Lagnado, 2019). These signals would allow hair cells activated by anterior deflection to encode the strength of a preceding swim episode.

Of the hair cells activated by anterior deflection, about 50% were unaffected and the other 50% only partially suppressed. A key question, therefore, is whether this population allow the lateral line to continue signaling mechanical stimuli while the fish is swimming; behavioural experiments by Feitl et al., (2010) indicate that the answer is yes. Using a suction source to mimic a predator they presented stimuli perpendicular to the body and compared escape responses in fish at rest and during swimming: an escape response was still triggered by a suction source while the larvae were swimming, although the probability decreased from 80% at rest to 40% when swimming. These behavioural observations are consistent with our neurophysiological results and indicate that suppression of self-generated signals does not come at the cost of losing all sensitivity to stimuli originating in the external world.

## STAR Methods

### CONTACT FOR REAGENT AND RESOURCE SHARING

Further information and requests for resources and reagents should be directed to and will be fulfilled by the Lead Contact, Leon Lagnado.

### EXPERIMENTAL MODEL AND SUBJECT DETAILS

#### Zebrafish

All procedures were in accordance with the UK Animal Act 1986 and were approved by the Home Office and the University of Sussex Ethical Review Committee.

Three zebrafish lines were used in this study. (1) The Tg[HuC::GCaMP6f] expresses GCaMP6f in all neurons (except for a small number of neuronal sub-types, including hair cells) and was kindly provided by Dr Isaac Bianco. (2) The Tg[Sill2, UAS::iGluSnFR] expresses the glutamate sensor iGluSnFR (Marvin et al 2013) under the control of the SILL promoter, which specifically targets afferent neurons of the posterior and anterior lateral line (Pujol-Marti et al 2012) and allows to measure hair cell glutamate release onto the afferents in the neuromast, as well as glutamate release by the afferent neurons in the hhindbrain. The generation of this line is described in Pichler & Lagnado (2019). (3) The (Tg[Sill2, UAS::iGluSnFR], cacnb^ts25/ts25^), is the same as (2) only in the background of the *‘relaxed’* (cacnb^ts25/ts25^) mutation (Granato et al 1996, Schredelseker et al 2005), which yields immotile homozygotes, due to a point-mutation in the b1a subunit of the dihydropyridine receptor involved in excitation-contraction coupling of skeletal muscle. It was generated by co-injecting the Sill2 and the 10xUAS::iGluSnFR plasmids (12 ng/μl) as well as the Tol2 transposase (40 ng/μl) (Kawakami 2007) into one-cell stage embryos originating from an in-cross of heterozygous *relaxed* mutants (cacnb^ts25/+^). Larvae were screened for expression of the iGluSnFR transgene and reared to adulthood. Founder fish, heterozygous for the *relaxed* mutation and carrying the Sill2 and iGluSnFR transgenes in their germline, were identified by outcrossing to heterozygous *relaxed* fish and screening the offspring for immobility (only the homozygotes are immotile) as well as the expression of iGluSnFR in lateral line afferents. As the homozygous *relaxed* larvae are not viable and die at 5 – 6 days post fertilization (dpf), the line was maintained in a heterozygous background and in-crossed to yield homozygotes, necessary for experiments.

Adult zebrafish were maintained in fish water at 28.5°C under a 14:10 hour light:dark cycle under standard conditions (Brand et al 2002). Fish were bred naturally, and fertilized eggs were collected, washed with distilled water and transferred into 50 ml of E2 medium (concentrations in mM: 0.05 Na2HPO4, 1 MgSO4 7H2O, 0.15 KH2PO4, 0.5 KCl, 15 NaCl, 1 CaCl2, 0.7 NaHCO3, pH7-7.5). At 24 hours post fertilisation (hpf) 1-phenyl2-thiourea (pTU) was added to yield a final concentration of 0.2 mM to inhibit pigment formation.

### METHODS DETAILS

#### Sample preparation

Sample preparation differed slightly between larvae in the wild-type background (Tg[HuC::GCaMP6f], Tg[Sill2, UAS::iGluSnFR]) and those in the immotile *relaxed* background (Tg[Sill2, UAS::iGluSnFR], cacnb^ts25/ts25^). Larvae of either sex were used in all experiments. The former were prepared as described earlier (Pichler & Lagnado, 2019), In brief, experiments were performed between 7-9 dpf, on larvae that were screened for the strongest expression of the respective transgene. They were anaesthetized in in 0.016% tricaine (MS-222) and were placed ‘side-down’ in a ‘fish-shaped’ pit, carved out of a thin layer of PDMS (Sylgard184, Dow Crowning) on a coverslip and held down by a ‘harp’ (Warner Instruments). Pressure of the Nylon strings was adjusted so that blood circulation was not compromised. Then, 0.25 mM a-Bungarotoxin (Tocris Bioscience) was injected into the heart to induce paralysis. Special care was taken to not touch the upward facing side of the fish, to avoid damaging the cupula.

Experiments on homozygote *relaxed* larvae were performed at 5 dpf. They were identified by the absence of movement upon tactile stimulation at 2 dpf and subsequently screened for the strongest expression of the iGluSnR transgene at 4 dpf. Preparation was carried out as mentioned above, only without using tricaine or a-Bungarotoxin.

#### Two-Photon Imaging

Two-photon imaging was performed as previously described (Pichler & Lagnado 2019). In short, a custom built two-photon microscope driven by a mode-locked Titanium-sapphire laser (Chameleon 2, Coherent) tuned to 915 nm (Odermatt et al 2012) was used. In experiments on larvae of the wild-type background excitation was delivered through a 40x water immersion objective (Olympus, 40x LUMIPlanF, NA: 0.8) and in experiments on the *relaxed* larvae, a 25 x objective (Nikon N25X-APO-MP 1.1NA) was used. To improve the signal-to-noise emitted photons were collected through the objective as well as through an oil condenser (NA 1.4, Olympus), below the sample. Green emission filters (525/70 nm at the objective and 530/60 nm at the condensor) were used in front of GaAsP photodetectors (H10770PA-40, Hamamatsu). The photocurrents of the two detectors were summed and passed through a transimpedance amplifier (Model SR570, Stanford Research Systems) and low-pass filtered (300 kHz). The microscope was controlled by ScanImage v3.8 (Pologruto et al 2003), synchronised with the stimulus application and operated at acquisition rates of 20-50 Hz. In this study, only neuromasts from the posterior lateral line (L3 – L6) with a directional sensitivity along the anterior-posterior axis were examined.

#### Mechanical stimulation

Mechanical stimulation was performed as described earlier (Pichler & Lagnado, 2019). Briefly, neuromasts were stimulated with positive and negative pressure steps, applied through a glass pipette (GC150T-10, Harvard Apparatus) with a tip diameter of ~30 μm, attached to a high-speed pressure clamp (HSPC-1, ALA scientific) (Trapani et al 2009). Output pressure was controlled through mafPC (courtesy of M. A. Xu-Friedman) running on IgorPro (Wavemetrics), which also triggered acquisition in ScanImage via a TTL pulse. The pipette tip, which was bent through ~30° using a micro forge (Narishige) to stimulate the neuromast approximately parallel to the body surface of the fish, was positioned ~ 20 μm above the body and ~100 μm away from the neuromast. The pressure clamp was manually zeroed before the start of an experiments so that no net flow was produced. We chose stimulus strengths that elicited near saturating responses in hair cells, assessed by a coarse protocol consisting of three positive and negative pressure steps of increasing amplitude (Pichler and Lagnado, 2019). Furthermore, the direction of the pipette relative to the fish (pointing anteriorly or posteriorly) was changed during the course of the experiments and no difference was observed.

#### Visual stimulation

In some experiments, we engaged the optomotor response by projecting a moving grating directly onto the larva, moving in the tail to head direction. A microprojector (Pico PK320, Optoma) from which the blue and green LED channels were removed was used to project a grating (12 mm wide bars at 100% contrast that moved at 5 mm/s) at an intensity that did not lead to bleed-through in the PMTs. The visual stimulus was controlled via the PsychoPy toolbox running in Python 3.6 and synchronized to the mafPC, controlling the mechanical stimulus, via a TTL pulse. In the set of experiments on the *relaxed* larvae, in which the 20x objective was used, visual stimulation was not as efficient in triggering fictive locomotion. This is most likely due to the objective being significantly wider and therefor restricted the light from the projector that actually reached the larvae.

#### Motor nerve recordings

Motor nerve recordings were performed as described by Masino & Fetcho (2005) with only minor modifications. Recording electrodes were pulled to a tip diameter of ~30 μm (from borosilicate glass, GC150T-10, Harvard Apparatus) and subsequently fire polished using a micro forge (Narishige). It was filled with extracellular recording solution (concentrations in mM: 134 NaCl, 2.9 KCl, 1.2 MgCl2, 2.1 CaCl2, 10 HEPES buffer, adjusted to pH 7.8 with NaOH). The pipette was positioned dorsally of the larva, above myotomal cleft 8-14 at a 45° angle and perpendicular to the longitudinal body axis. Using a plastic syringe, slight positive pressure was applied during the approach and upon contacting the skin changed to negative pressures between −30 and −70 mmHg. On average spontaneous motor nerve activity could be observed after 10-15 minutes. Using a BVC-700A (Dagan, USA) in current-clamp mode the extracellular voltage was measured. The signal was filtered (Brownlee model 440, Neurophase) with a high and low-pass cut off frequency of 300 Hz and 1 kHz, respectively and recorded using mafPC at a sample rate of 5 kHz (synchronously with the mechanical stimulation

### QUANTIFICATION AND STATISTICAL ANALYSIS

#### Image segmentation and analysis

Images sequences (movies) were analyzed in Igor Pro. Small drifts in the x/y dimension were registered, using the SARFIA toolbox (Dorostkar et al 2010). Movies with large drifts, and potential z-drifts were discarded. Regions of Interest (ROIs) were determined using a custom written procedure (analogous to Portugues et al 2014, and described in Johnson et al 2019) that identifies pixels with the highest correlation value (to neighboring pixels) as ‘seeds’ and extends these to form ROIs, based on a threshold, manually defined by the experimenter. These ROIs corresponded to sites of maximal glutamate release which occur in apposition to hair cell ribbon synapses (Pichler & Lagnado, 2019).

Background fluorescence was subtracted manually. Baseline fluorescence (F) was defined as the average fluorescence in the first 10 s of imaging and preceding the first stimulation interval; the ratio of change in fluorescence (ΔF) was calculated relative to that value (ΔF/F) and used for further analysis. Contrast in images was adjusted for presentation purposes. The motor nerve recordings were further digitally filtered (300 Hz high-pass 1kHz low-pass and 50 Hz notch). Spikes were extracted using a custom written procedure that applied a simple threshold to the filtered signal and detected when it was crossed by the signal. This temporal filter was a Gaussian with FWHM = 100 ms.

---

**Figure S1.**
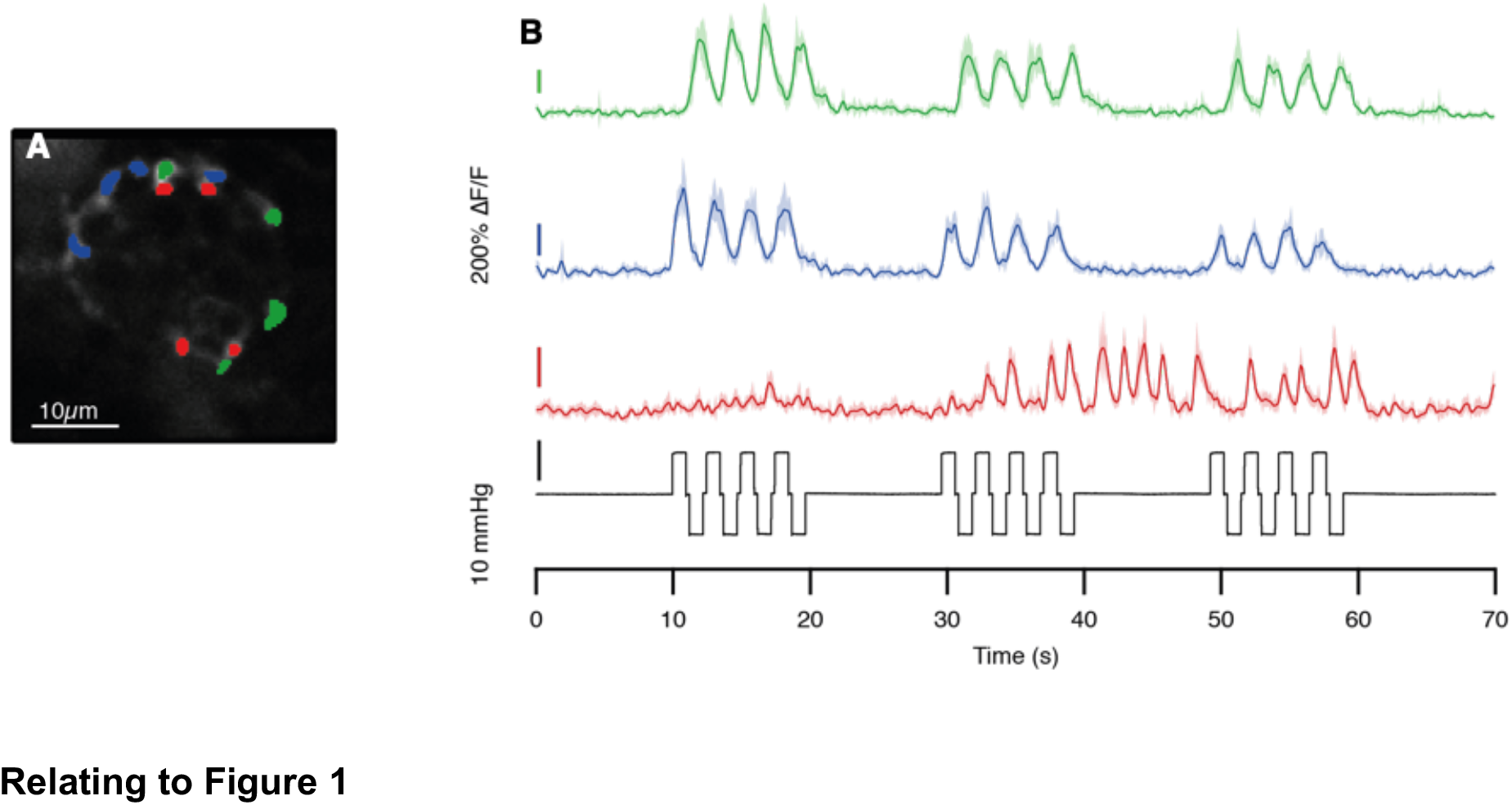
Efferent synapses can be identified based on their morphology and firing pattern, which is independent of mechanical stimulation. (**A**) Zebrafish larvae expressing the calcium indicator GCaMP6f under the control of the HuC promoter (Tg[HuC::GCamP6f]) were paralyzed with α-BTX. An average projection of a NM, highlighting irregularly shaped varicosities belonging to two afferents with opposing directional sensitivity (blue and green) as well as small and round efferent boutons (red). (**B**) The average signals from the respective ROIs in (A). Blue ROIs are sensitive to posterior deflections and green ROIs to anterior, confirming that they are afferents. Red ROIs fired independent of the mechanical stimulation, confirming that they were efferent. Shaded area represents SEM.

**Figure S2.**
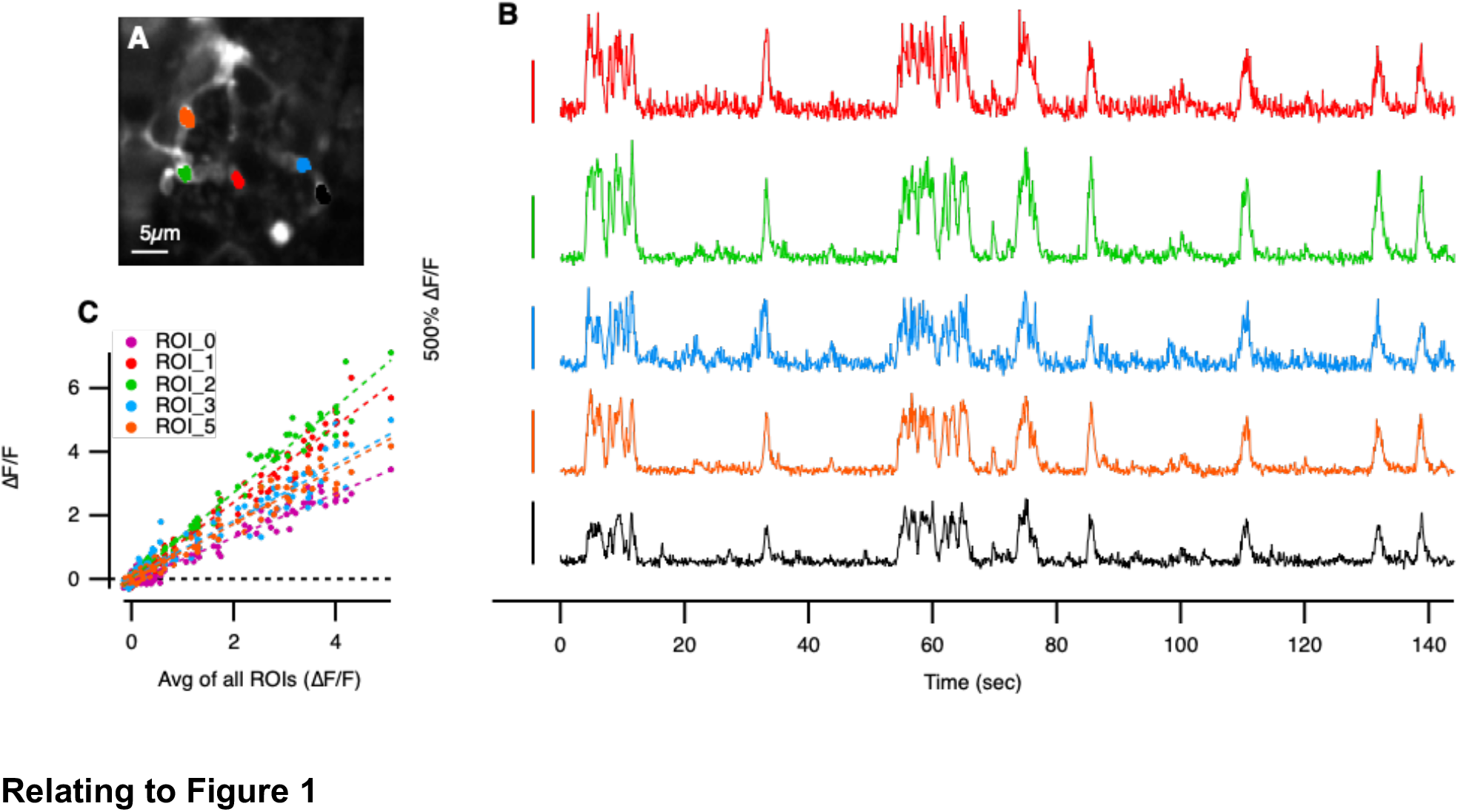
Activity is strongly synchronized across efferent synapses. (**A**) Zebrafish larvae expressing the calcium indicator GCaMP6f under the control of the HuC promoter (Tg[HuC::GCamP6f]) were paralyzed with α-BTX. Efferent ROIs were identified based on their, small, roundish morphology as well as their ‘spontaneous’ activity in the absence of mechanical stimulation. (**B**) Response profile of the 5 ROIs depicted in (A) over 145 s time window. These were all “spontaneously” activity in the absence of mechanical stimulation. (**C**) Plot of the instantaneous signal in each of the five synapses as a function of the average activity of all five. Activity in B was down-sampled into time bins of 2 s. Each set of data points could be fit by a straight line through the origin with r > 0.9 revealing a high degree of synchronicity.

**Figure S3.**
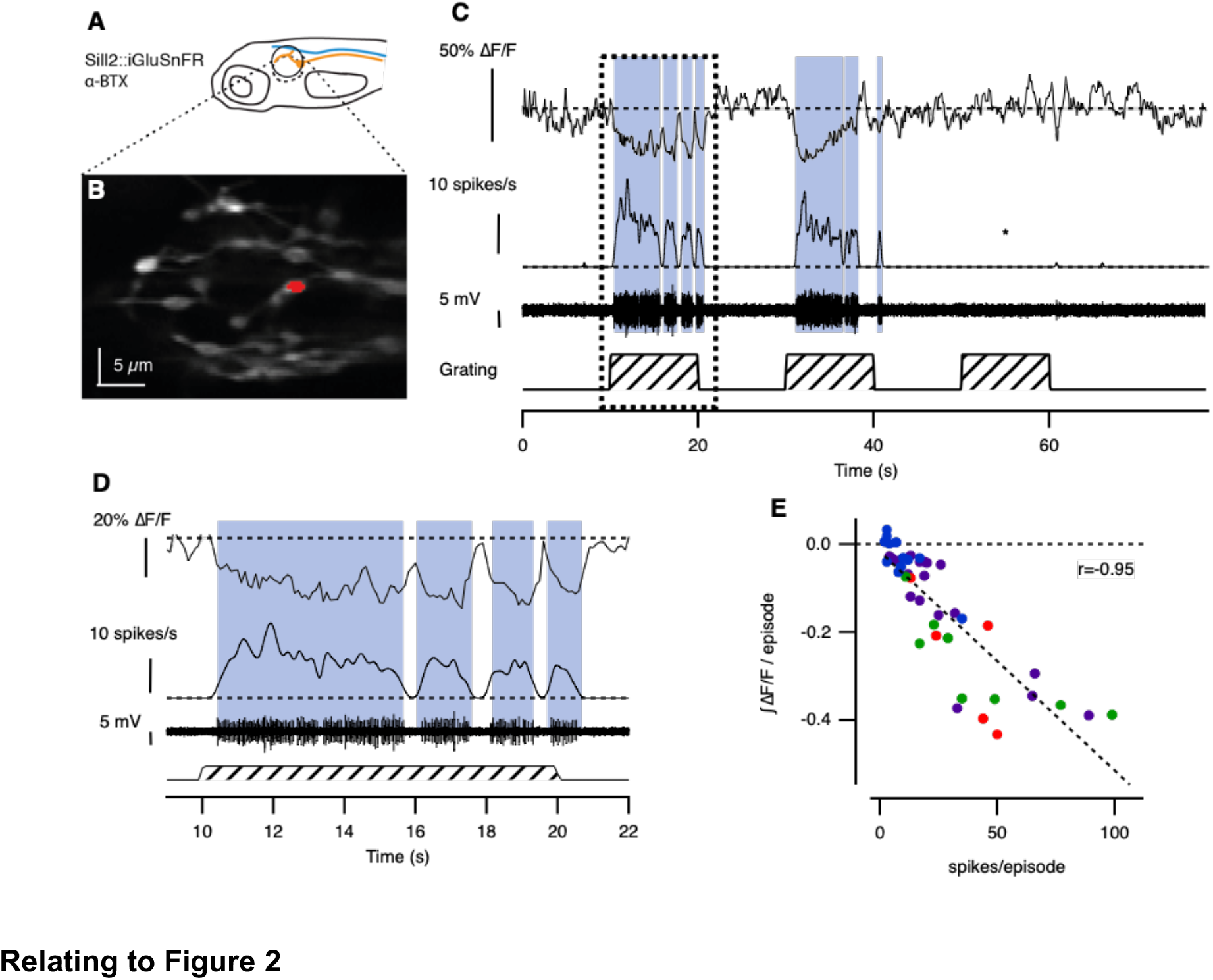
Motor activity suppressed spontaneous activity in afferent fiber synapses transmitting to the medial octavolateralis nucleus (MON). (**A**) Experiments were carried out in larvae that expressed the glutamate reporter iGluSnFR in afferent neurons (Tg[Sill2, UAS::iGluSnFR] and which were paralyzed with α-BTX. (**B**) An average projection of the posterior arm of the MON, highlighting the synapse whose response is depicted in (C). (**C**) An example of a synapse, whose baseline glutamate release, in the absence of a mechanical stimulus, was suppressed by fictive swimming (grey-shaded area). The absence of suppression during the third presentation of a visual grating (*) indicated that the visual stimulus did not directly affect the encoding of mechanical information. (**D**) Magnification of the dashed box in (C) reveals that each individual swim-burst leads to a transient suppression of the glutamate release. (**E**) The number of spikes in the motor neuron during a swim-bout (episode) and the integral of the suppressive effect in the hindbrain synapse is tightly correlated (r=−0.95, n=49 bouts from 4 synapses).

**Figure S4.**
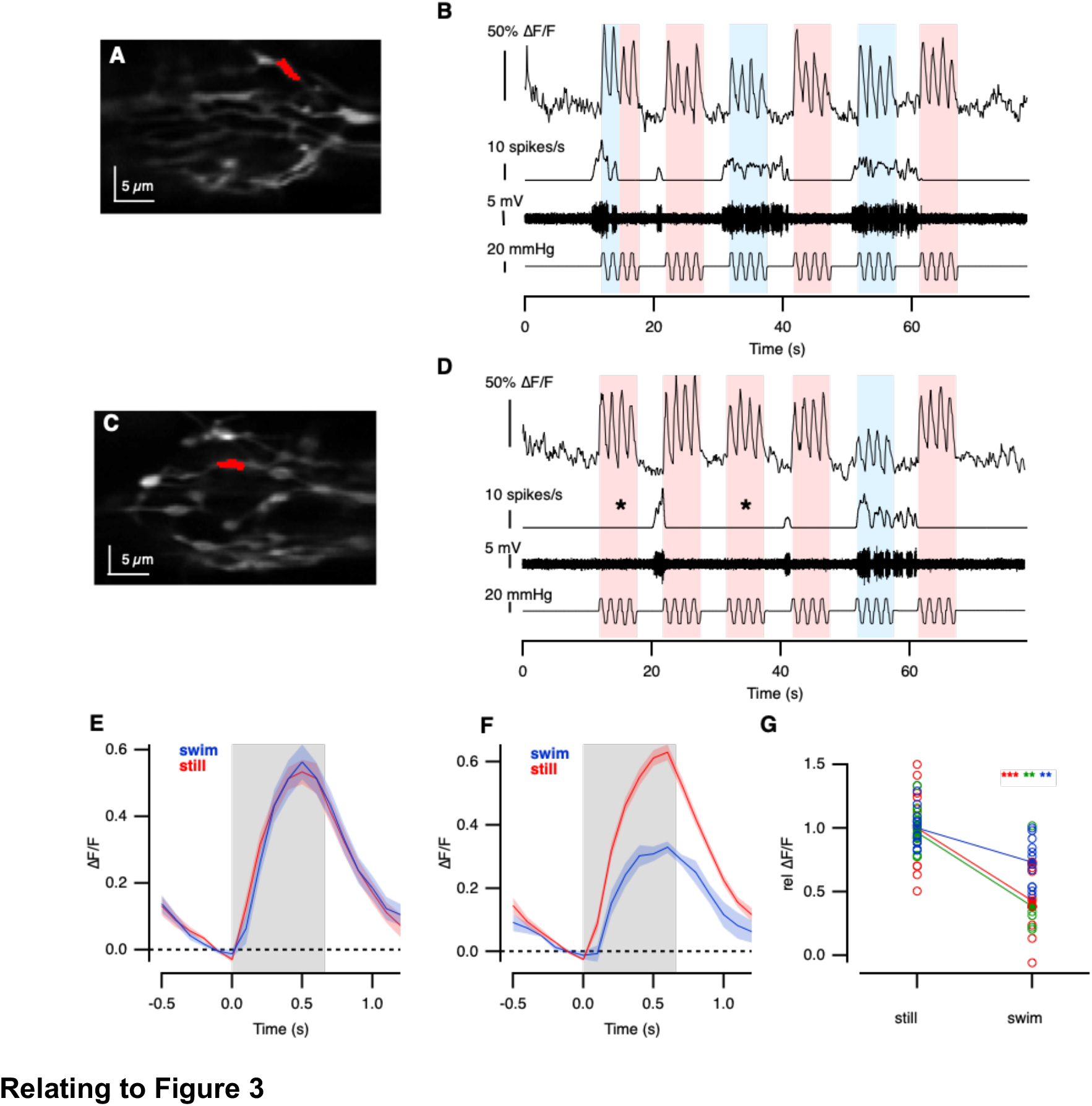
Motor activity suppressed stimulus-evoked activation of afferent fiber synapses transmitting to the medial octavolateralis nucleus (MON) in the hindbrain. Experiments were carried out in larvae that expressed the glutamate reporter iGluSnFR in afferent neurons (Tg[Sill::Gal4, UAS::iGluSnFR] and which were paralyzed with α-BTX. A given neuromast was stimulated with positive and negative pressure steps while the output of afferent neurons in the MON was imaged with a two-photon microscope. (**A** and **C**) Average projections of synapses in the posterior part of the MON. Highlighted in (A) is the synapse whose response is depicted in (C) and highlighted in (C) is the synapse whose response is depicted in (D). (**B** and **D**) Two representative examples of afferent synapses in the MON, which are sensitive to posterior deflections. The bottom trace represents the mechanical stimulation of the NM (positive pressure corresponds to posterior and negative pressure steps to anterior deflection of the cupula, respectively). The middle two traces depict the raw motor signal and average spike rate, respectively, and the top trace shows the synaptic activity of the synapse. Blue-shaded areas indicate periods in which mechanical stimulation coincided with fictive swimming and red-shaded areas indicate periods in which it did not. The first third and fifth mechanical stimulation period always overlapped with the presentation of a moving grating to induce fictive locomotion. (**B**) The activity of this synapse was not affected by fictive locomotion. This is quantified in (E), which depicts the mean responses of this synapse in the presence and absence of fictive locomotion. (**D**) Example of a synapse whose response to mechanical stimulation was suppressed when it coincided with fictive locomotion, quantified in (F). Asterisks indicate periods in which visual stimulation failed to induce locomotion. (**F**) The peak amplitude of the IGluSnFR signal in the MON was reduced by 42% during fictive locomotion (P<0.005, Mann-Whitney U-test). (**G**) The mechanically induced iGluSnFR signal in three hindbrain synapses while still (left) and during ‘fictive swimming’ (right). All these afferents were activated by posterior deflections of the cupula. Open circles represent the response to individual stimulations and filled circles their average (*** P<0.0001, ** P<0.001, Mann-Whitney U-test). (Shaded areas in (E) and (F) represent the SEM).

